# LingoCell interprets gene perturbations through natural language and tissue niches

**DOI:** 10.64898/2026.11.08.765157

**Authors:** Yitao Yang, Wanzhe Xu, Zhenhua Zhang, Yang Cui, Xin Yu, Wenchao Gu, Junjian Wang, Kenta Nakai

## Abstract

Single-cell foundation models are general tools for resolving cell identities and disease states, but struggle to interpret changes in expression magnitude in biological language, to transfer to sparse spatial measurements, and to explain cellular niches. We introduce LingoCell, a foundation model connecting single-cell and spatial transcriptomics. It aligns cell representations with language while keeping gene expression editable, supporting cell annotation, cross-batch integration, description generation, and disease-language readouts of single-gene and combinatorial *in silico* perturbations. Extended to tissues, it annotates cells without single-cell references, learns identity-conditioned niche embeddings, and performs spatial *in silico* perturbation by replacing and editing neighbor expression. In lung adenocarcinoma, LingoCell revealed synergistic disease responses to jointly weakening alveolar differentiation programs and strengthening malignant programs. In colorectal cancer, it revealed different programs conveyed by the same fibroblast state to T cells, macrophages, and tumor cells. LingoCell provides a unified framework for prioritizing target combinations and stromal intervention programs.

## Introduction

Single-cell foundation models learn gene coordination by pretraining on tens of millions of cells to support cell annotation, batch integration, and perturbation analysis, yet their perturbation and interpretation capabilities remain constrained by how expression is represented. Geneformer represents cells as gene sequences ranked by relative expression and identified cardiomyopathy candidate targets through *in silico* deletion,^1^ but rank-based encoding records only relative gene order, preventing magnitude adjustments that preserve rank order from entering the model. scGPT jointly models gene identity and discretized expression values to support annotation, integration, and fine-tuned perturbation prediction,^2^ while CellFM scaled pretraining to roughly 100 million human cells.^3^ Although these models detect magnitude changes, their perturbation outputs are embedding shifts or predicted expression profiles rather than semantic readouts defined by disease descriptions. More fundamentally, none is aligned during pretraining with natural language describing cell identity, function, and disease, making existing biological concepts difficult to use for identifying cells in new data or interpreting molecular changes.

Aligning cell representations with language brings these concepts directly into the model. LangCell achieves zero-shot annotation through cell–text contrastive learning but retains rank-based encoding and does not generate cell descriptions.^4^ CellWhisperer links transcriptome–text alignment to a large language model for free-text cell retrieval and querying, but its analyses stop at observed expression.^5^ Cell2Text generates cell descriptions from frozen Geneformer features, but whether expression changes map to disease-semantic responses remains untested.^6^ Interpreting gene changes through disease language requires retaining editable expression magnitude while aligning with text.

Even when a model interprets and edits a cell’s own expression through language, models trained on dissociated cells lack cellular position, adjacency, and morphology in tissues. Platforms such as Visium HD have advanced spatial transcriptomics to near-single-cell resolution.^7^ Methods such as STAGATE and GraphST integrate expression and spatial adjacency to identify tissue domains,^8,9^ but require training on target data, do not directly output cell identities, and typically need a matched single-cell reference. scGPT-spatial extends continual pretraining across spatial platforms,^10^ yet spatial capture depth is far lower than in single-cell sequencing and expression distributions of the same cell type differ systematically; adapting to spatial distributions alone does not guarantee preservation of single-cell identity knowledge, leaving open how to sustain cell annotation and semantic understanding in spatial data.

Maintaining stable cell identities in tissues is also fundamental to understanding niches: the same environment conveys different information to cells with different functions, and neighbor features relevant to a T cell need not be those to which a tumor cell responds. Existing methods mainly define niches from the environment itself: BANKSY combines a cell’s own expression, neighborhood mean expression, and spatial gradients,^11^ whereas DECIPHER uses cross-scale contrastive learning to represent molecular identity and spatial environment separately.^12^ Neither environmental representation is queried by the central cell’s identity, making environmental differences difficult to interpret in a specific cellular context. Interpreting niche differences must ultimately resolve neighbor gene programs, yet localization-associated gene analysis yields only observational associations and cannot answer how altering a neighbor program changes the environment of a specific cell.

We therefore introduce LingoCell, a foundation model connecting cell identity, disease concepts, and tissue environment through language-aligned expression representations. Pretrained on more than 11 million single cells and their paired descriptions, LingoCell jointly encodes gene identity and expression level and aligns them with text, supporting clustering, cross-batch integration, zero-shot annotation, and description generation without fine-tuning; because expression magnitude remains editable after alignment, it can also adjust one or more genes and read cellular responses using interchangeable disease descriptions. Transferred to tissues through measurement adaptation, LingoCell annotates Visium HD data at single-cell resolution without a single-cell reference, uses the central cell’s identity as a query to build observer-dependent niche embeddings integrating continuous neighbor molecular states and spatial structure, and performs spatial *in silico* perturbation by replacing and editing neighbor states. Using LingoCell, we found that co-downregulating alveolar differentiation programs and co-upregulating malignant programs synergistically increased cellular alignment with disease descriptions in lung adenocarcinoma, distinguished fibroblast environments with similar neighbor compositions but different molecular states in normal colon, and revealed distinct program priorities for T cells, macrophages, and tumor cells reading the same fibroblast state in colorectal cancer. As a unified foundation model spanning cell atlases and tissues in situ, LingoCell enables researchers to investigate the meaning of genes or neighbor programs across single-cell and tissue scales, advancing analysis from describing what cells are to interpreting what molecular changes mean.

## Results

### Overview of LingoCell

LingoCell connects single-cell and spatial transcriptomics within a unified architecture (Fig. 1): the single-cell stage aligns editable gene expression with natural language to support representation analysis, description generation, and disease-semantic perturbation (Fig. 1a–d), whereas the spatial stage transfers to tissues in situ through measurement adaptation to resolve identity-conditioned local environments and perform neighborhood *in silico* perturbation (Fig. 1e–g).

**Fig. 1.**
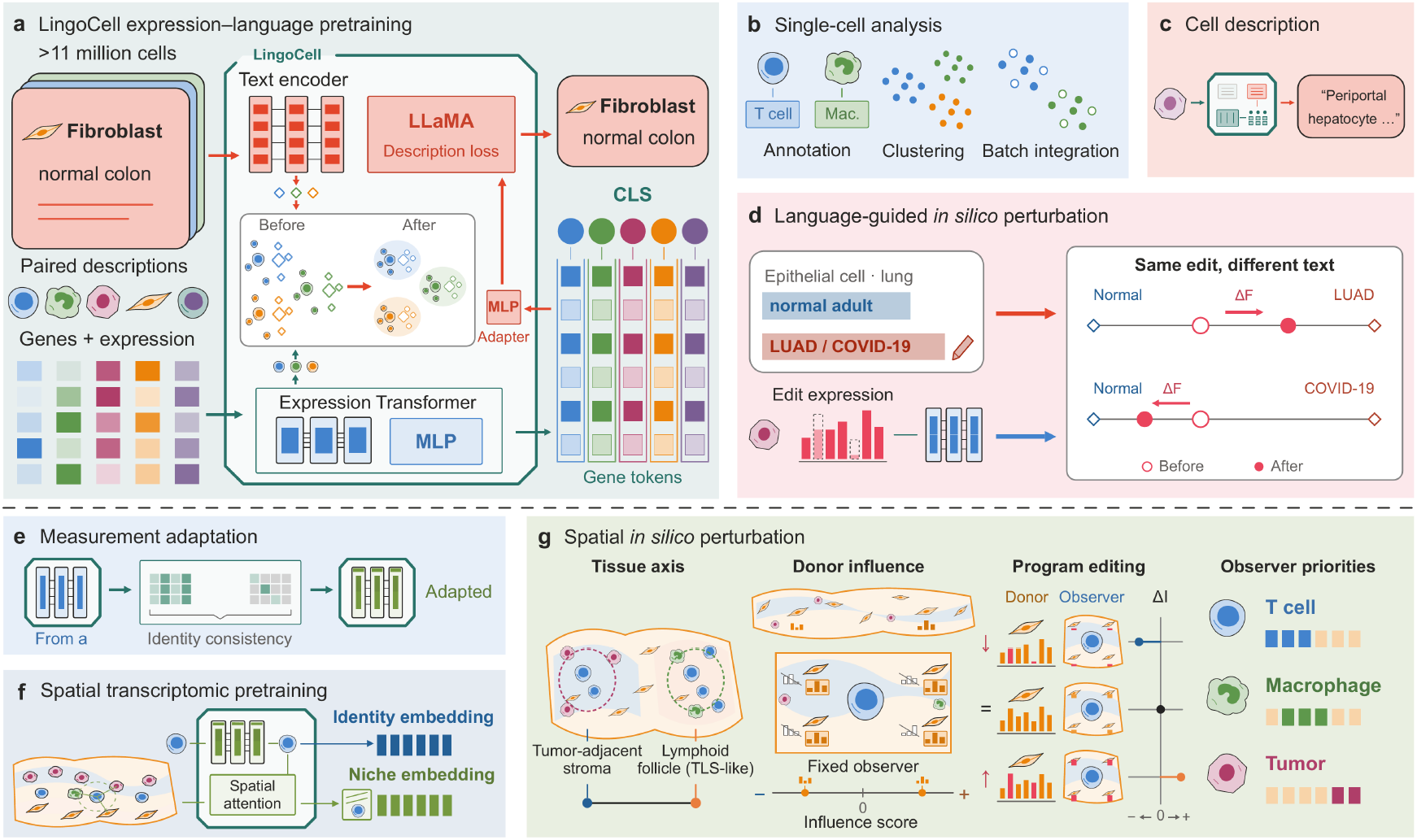
**LingoCell framework connecting single-cell expression, natural language, and tissue neighborhoods**. **a,** Expression–language pretraining: gene identities, expression levels, and paired descriptions of single cells are encoded by the expression Transformer and text encoder, respectively. Expression recovery, cross-modal alignment, and cell-type prototype learning form a shared representation space, and gene features are passed through an adapter to the LLaMA decoder to generate cell descriptions. **b,** Identity embeddings are used for cell annotation, clustering, and cross-batch integration. **c,** Cell descriptions are generated from expression. **d,** Language-guided single-cell *in silico* perturbation: normal and disease descriptions define a semantic direction, cells are re-encoded after gene expression levels are adjusted, and the semantic shift along this direction (ΔF) measures the response; the disease reference is interchangeable. **e,** Spatial measurement adaptation: changes in measurement depth and sparsity are simulated to adapt the expression encoder to spatial measurements through objectives including identity consistency. **f,** Spatial pretraining: central-cell expression, neighbor expression, spatial distances, and H&E images are integrated to produce identity embeddings and niche embeddings referenced to identity. **g,** Spatial *in silico* perturbation: tumor-adjacent stroma and lymphoid follicle (TLS-like) environments define a niche reference axis, states of real fibroblast donors are inserted into observer neighborhoods, axial shifts of observer niches measure donor influence, and differences in influence before and after gene program upregulation or downregulation (ΔI) compare the program emphasis of different observers. CLS, identity features summarizing cell information; gene tokens, gene features; TLS, tertiary lymphoid structure.

Single-cell pretraining uses more than 11 million single cells and paired descriptions from CELLxGENE (Fig. 1a).^13^ The expression encoder reads gene identity and expression level, while the text encoder reads cell-type, tissue, and state descriptions; cross-modal contrastive learning and cell-type prototype learning bring both into a shared representation space, and expression recovery preserves gene-level information. Frozen identity embeddings directly support annotation, clustering, and cross-batch integration (Fig. 1b), while gene representations pass through an adapter to a LLaMA decoder to generate natural language describing cell identity, tissue origin, and function (Fig. 1c). Because expression magnitude remains editable, adjusting one or more genes in a fixed cellular context and re-encoding yields a semantic shift (ΔF) along the direction defined by normal and disease descriptions as a disease-semantic readout; switching the disease description without changing the perturbation interprets the same expression change against different diseases (Fig. 1d).

On entering the spatial stage, because platforms such as Visium HD capture expression more sparsely at single-cell resolution and differ systematically from single-cell sequencing, LingoCell first performs measurement adaptation training on single-cell data, simulating spatial capture depth and sparsity while constraining the same cell to retain consistent identity embeddings across observation conditions (Fig. 1e). The adapted encoder then joins a spatial attention module for self-supervised pretraining on real tissue sections (Fig. 1f and Supplementary Table S1): using the central cell’s identity as a query, the spatial module reads neighbor expression, distances, and tissue morphology, learns tissue context through expression recovery and prediction of neighborhood cell composition and pathway activity, and outputs an identity embedding for spatial annotation and a niche embedding referenced to that identity.

Niche embeddings extend *in silico* perturbation to tissue neighborhoods (Fig. 1g): fixing the central cell (observer) and its spatial context, real states of a given neighbor type from across the section (donors) are placed into its neighborhood one at a time, and the shift of the observer’s niche embedding along a reference axis defined by two real tissue environments measures donor influence; candidate programs built from influence-associated donor genes are then upregulated or downregulated *in silico*, and the resulting influence difference (ΔI) tests effect direction and compares how different observers weight the same neighbor state.

### LingoCell organizes single-cell representations around cell-type prototypes for clustering, integration, and zero-shot annotation

We evaluated frozen pretrained LingoCell representations without fine-tuning on four public datasets: PBMC10K (peripheral blood mononuclear cells), cross-platform Pancreas, Human Immune (blood and bone marrow), and multi-organ Tabula Sapiens.^14–16^ Comparators included expression foundation models (scGPT, Geneformer, CellFM), language-aligned models (LangCell, CellWhisperer), and standard representations (PCA and highly variable genes, HVG), with clustering and integration metrics computed under the scIB framework.^15^

In the pretraining atlas, LingoCell organized representations around cell identity, separating 24 displayed categories in a representative subset (Fig. 2a). Across the four benchmark datasets, LingoCell achieved the highest mean Leiden clustering adjusted Rand index (ARI, 0.584) and normalized mutual information (NMI, 0.742) among the eight representation methods, outperforming the runner-up CellWhisperer (mean ARI 0.517) and PCA (mean NMI 0.733), with top ARI on every dataset (Fig. 2b, Supplementary Fig. S1, Supplementary Table S2, and Supplementary Note 1). In PBMC10K, LingoCell separated CD14⁺ monocytes, FCGR3A⁺ monocytes, and dendritic cells into distinct adjacent regions, whereas CellWhisperer showed more intermingled boundaries (LingoCell ARI 0.887 and NMI 0.843, versus runner-up CellWhisperer 0.828 and PCA 0.832; Fig. 2c).

**Fig. 2.**
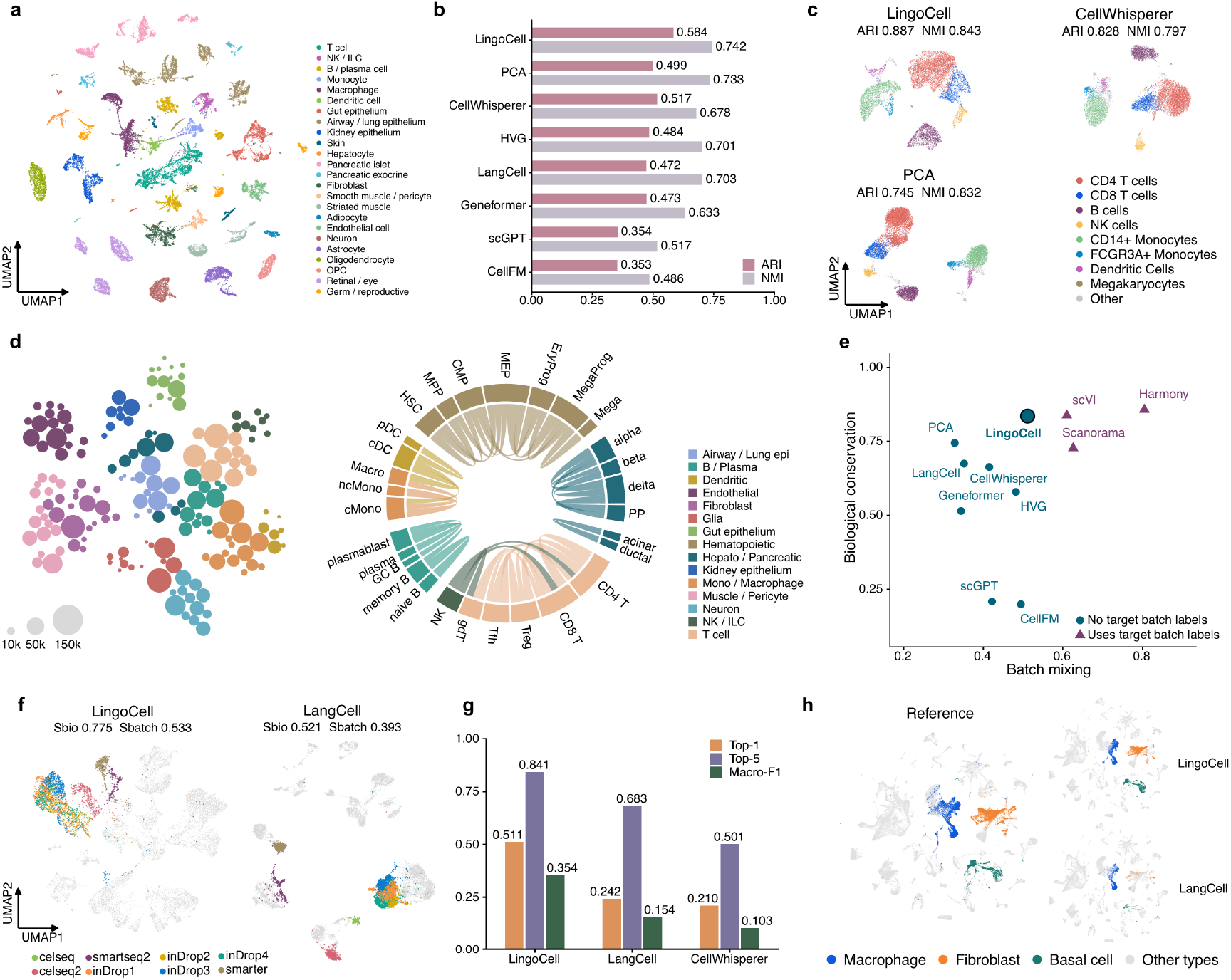
**Identity embeddings, prototype organization, and basic single-cell functions of LingoCell**. **a,** Identity embedding UMAP of representative cells in the pretraining atlas, colored by cell class. **b,** Mean Leiden ARI and NMI of eight representation methods across four benchmark datasets. **c,** UMAPs of LingoCell, CellWhisperer, and PCA on PBMC10K, colored by reference cell type and annotated with ARI and NMI. **d,** Cell-type prototype landscape (left; circle area is proportional to the number of pretraining cells of that type) and a chord diagram of cosine similarities between representative prototypes (right; only connections with similarity ≥0.30 are shown). **e,** Mean biological conservation (Sbio) and batch mixing (Sbatch) scores across Pancreas, Human Immune, and Tabula Sapiens; triangles indicate integration methods that use target batch labels. **f,** UMAPs of LingoCell and LangCell on Pancreas, with β cells colored by batch and other cells gray, annotated with Sbio and Sbatch. **g,** Top-1 accuracy, Top-5 accuracy, and macro-F1 for zero-shot annotation by three language-aligned models on Tabula Sapiens. **h,** UMAPs of reference annotations and LingoCell and LangCell predictions on Tabula Sapiens, highlighting macrophages, fibroblasts, and basal cells. ARI, adjusted Rand index; NMI, normalized mutual information; macro-F1, macro-averaged F1.

This clustering advantage reflects the cell-type prototypes introduced during pretraining: expression representations and descriptions of the same cell type are gathered around their type prototype, drawing each cell toward a shared identity reference. In the prototype landscape, lymphoid, myeloid, epithelial, stromal, and neural families occupied concentrated regions, and strong connections in the similarity chord diagram occurred mainly within related families (Fig. 2d), indicating that the prototype bank also encodes lineage relationships above the cell-type level.

Identity-centered representations likewise favored cross-batch integration. Across Pancreas, Human Immune, and Tabula Sapiens, LingoCell achieved the highest mean biological conservation (0.853) and batch mixing (0.505) scores among the eight methods not using target batch labels (Fig. 2e); compared with Harmony, scVI, and Scanorama, which explicitly use batch labels,^17–19^ its biological conservation matched scVI (0.853 versus 0.855) and exceeded Scanorama, while its overall score on Tabula Sapiens (0.797) surpassed all three dedicated integration methods (Supplementary Fig. S2 and Supplementary Table S3). In cross-platform Pancreas, β cells from different batches converged into a single identity region in LingoCell but separated by sequencing protocol in LangCell (full-dataset biological conservation 0.775 versus 0.521, batch mixing 0.533 versus 0.393; Fig. 2f).

Language alignment further enables direct text querying of identity embeddings using descriptions of candidate cell types and tissue context without training a classifier. On Tabula Sapiens, LingoCell’s Top-1 accuracy (51.1%), Top-5 accuracy (84.1%), and macro-F1 (0.354) exceeded those of LangCell and CellWhisperer, more than doubling both on Top-1 and exceeding LangCell by 15.9 percentage points on Top-5 (Fig. 2g); on shared-layout UMAPs,^20^ its predictions for macrophages, fibroblasts, and basal cells better matched reference annotations (Fig. 2h). Zero-shot results on Human Immune and Pancreas and labeled-reference comparisons with scANVI and CellTypist are shown in Supplementary Fig. S3 and Supplementary Tables S4a, S4b, and S5,^21,22^ where LingoCell ranked in the top three of the eight representation methods in mean macro-F1.

### LingoCell translates single-cell expression profiles into molecularly grounded natural language descriptions

Beyond querying cells with given text, LingoCell can also generate open-ended descriptions de novo from transcriptomes: gene representations from the frozen expression encoder pass through an adapter to the language decoder, translating expression directly into coherent sentences summarizing cell identity, tissue origin, and function (Fig. 1a,c). Compared with Cell2Text, which generates descriptions from single-cell expression,^6^ LingoCell achieved RoBERTa- and BioBERT-based BERTScores of 96.60% and 94.20% (Cell2Text: 96.28% and 93.90%) and extracted cell-type, tissue, and disease F1 scores of 78.72%, 80.65%, and 79.50%, all exceeding Cell2Text (Fig. 3a), with consistent gains in field accuracy and lexical metrics including BLEU and ROUGE (Supplementary Fig. S4a,b and Supplementary Note 2). Generated text also links identity to biological function: kidney cells expressing AQP3, FXYD4, and SCNN1B were described as renal principal cells regulating sodium and potassium balance, and brain cells expressing PLP1, CNP, and MBP as oligodendrocytes forming axonal myelin in the central nervous system (Fig. 3b; additional examples in Supplementary Fig. S4c).

**Fig. 3.**
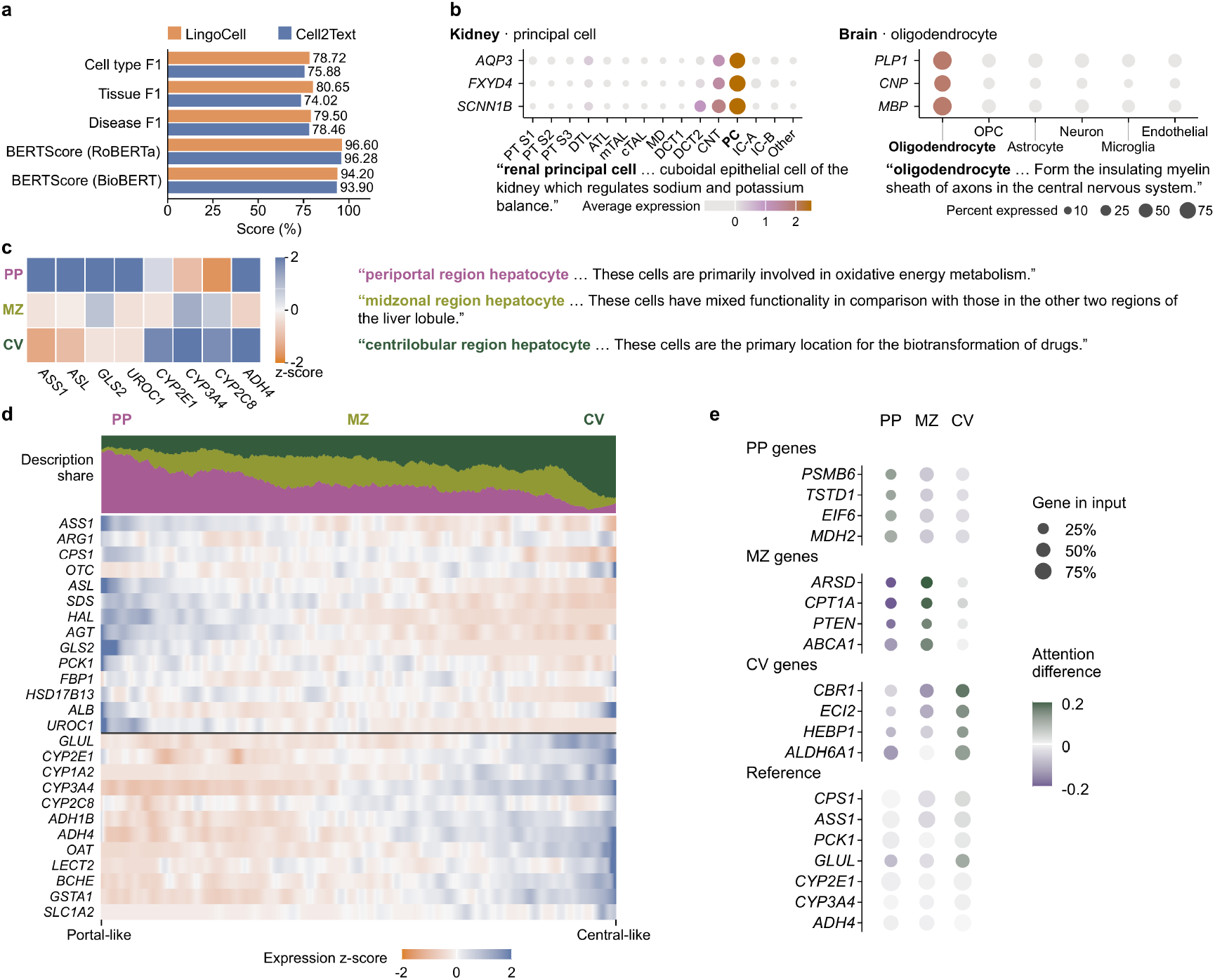
LingoCell generates natural language descriptions from single-cell expression and resolves continuous hepatocyte zonation. a, Weighted F1 for cell-type, tissue, and disease information in descriptions generated by LingoCell and Cell2Text (Llama-1B model), and BERTScore based on RoBERTa and BioBERT. **b,** Excerpts of generated descriptions for a renal principal cell and an oligodendrocyte, with generated identity phrases in bold; dot plots above show the mean expression (color) and proportion of expressing cells (dot size) for three marker genes across cell types in the same dataset, with the types of the example cells in bold. **c,** Expression z scores of zonation marker genes in three representative hepatocytes and corresponding excerpts of periportal, midzonal, and pericentral descriptions. **d,** Hepatocytes assigned zonation descriptions arranged along a periportal–pericentral expression coordinate, with local proportions of the three description classes above and smoothed expression z scores of zonation-related genes below. **e,** Cross-attention associations between zonal identity phrases and input genes, grouped into zone-associated genes and canonical reference genes; color indicates the difference in mean attention rank relative to other phrases, and dot size the proportion of cells in the group with the corresponding gene in the input. PP, periportal; MZ, midzonal; CV, pericentral.

Continuous state differences within a single cell type further test description resolution. Hepatocytes exhibit zonated metabolic specialization along the portal-to-central lobular axis.^23^ In the human liver single-cell data of MacParland et al.,^24^ nearly all selected hepatocytes received explicit zonation descriptions, with three representative cells summarized as periportal (oxidative energy metabolism), midzonal (mixed functionality), and pericentral (drug biotransformation), matching their zonation marker expression (Fig. 3c). Ordered along an expression coordinate defined by periportal (ASS1, HAL) and pericentral (GLUL, CYP2E1, CYP3A4) genes (Fig. 3d), or along a coordinate built from markers of the healthy human liver spatial atlas of Yakubovsky et al. (Supplementary Fig. S4d),^23^ generated descriptions shifted smoothly from predominantly periportal to predominantly pericentral. Cross-attention while generating identity phrases further showed that midzonal descriptions attended more to CPT1A and ABCA1, whereas pericentral descriptions attended more to CBR1 and ALDH6A1 (Fig. 3e), giving the text output a traceable molecular basis.

### LingoCell resolves disease-associated cell states and gene programs through semantic queries and *in silico* perturbation

The unified expression–language space enables free text to serve directly as a semantic probe of cellular function and disease states. In the lung single-nucleus atlas of Melms et al. (GSE171524), comprising COVID-19 decedents and controls,^25^ keeping identity embeddings fixed while querying “Angiogenesis” concentrated high-response cells in endothelial cells, switching to “Epithelial–mesenchymal transition” (EMT) shifted the response to mesenchymal fibroblasts, and “Complement system” highlighted immune populations including B cells, antigen-presenting cells, and T cells (Fig. 4a). These functional concepts map biological questions directly onto cell populations without prior conversion into marker gene lists.

**Fig. 4.**
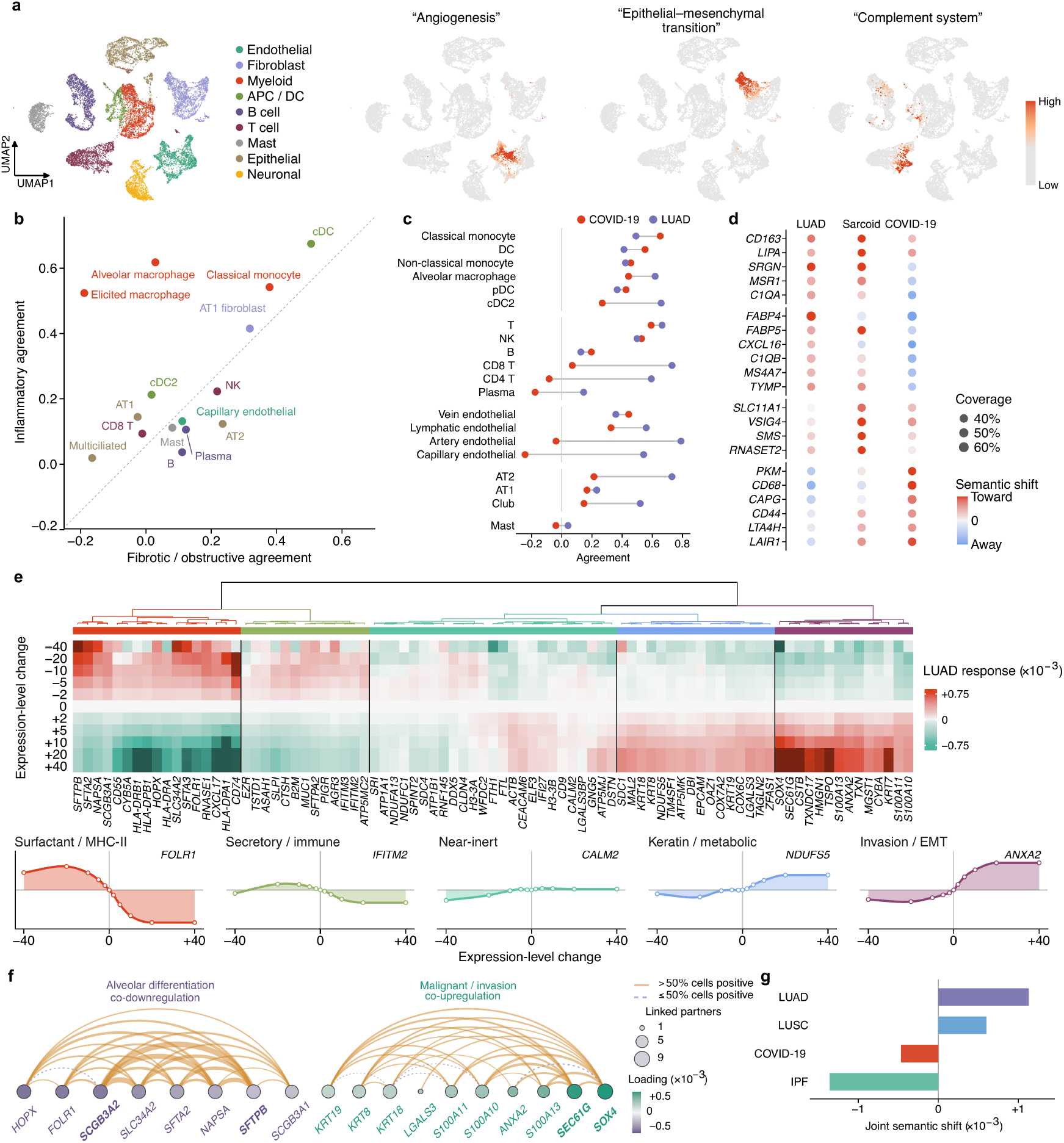
LingoCell links semantic queries to *in silico* perturbation of disease-associated gene programs. a, UMAP of the GSE171524 lung single-nucleus atlas, colored by cell type and scores for the queries “Angiogenesis”, “Epithelial–mesenchymal transition”, and “Complement system”. **b,** Alignment of normal-to-disease representation changes in lung cell types with language directions for inflammation-associated lung diseases (HP, NSIP, and pulmonary sarcoidosis) and fibrotic/obstructive lung diseases (PF and COPD). Colored as in a; the dashed line indicates equality. **c,** Alignment of normal-to-disease representation changes in lung cell types with COVID-19 and LUAD language directions, grouped by cell lineage. **d,** Mean semantic shifts along the LUAD, pulmonary sarcoidosis, and COVID-19 directions after upregulating candidate genes in healthy alveolar macrophages. Red, toward the corresponding disease description; blue, away from it; dot size, proportion of cells covered by each gene. **e,** Responses along the LUAD direction to different expression adjustment levels for each gene in LUAD epithelial cells, clustered into five groups by response pattern, with response curves of representative genes below. **f,** Two-gene connection diagrams for co-downregulation within the alveolar differentiation program (left) and co-upregulation within the malignancy/invasion-associated program (right). Arcs indicate positive non-additive effects (line width, effect size; solid and dashed lines, positive effects in more than half and no more than half of cells, respectively); node color indicates single-gene loading and node size the number of connected genes. **g,** Mean joint semantic shifts of the same cell states after NAPSA and SFTPB co-downregulation along the LUAD, LUSC, COVID-19, and IPF text directions. HP, hypersensitivity pneumonitis; NSIP, nonspecific interstitial pneumonia; PF, pulmonary fibrosis; COPD, chronic obstructive pulmonary disease; LUAD, lung adenocarcinoma; LUSC, lung squamous cell carcinoma; IPF, idiopathic pulmonary fibrosis.

Switching queries to disease descriptions distinguishes state changes across cell populations within an organ. Using CELLxGENE lung disease data, we compared normal-to-disease representation changes against language directions for inflammation-associated versus fibrotic/obstructive lung diseases (Fig. 4b): alveolar and elicited macrophages were biased toward the inflammatory side, conventional dendritic cells and classical monocytes responded strongly in both directions, and alveolar type II (AT2) cells were biased toward the fibrotic/obstructive side. Comparing COVID-19 and lung adenocarcinoma (LUAD) directions, the former highlighted classical monocytes, dendritic cells, and subsets of lymphocytes, whereas the latter involved immune, vascular endothelial, and alveolar epithelial cells, with capillary endothelial cells showing opposite alignment across the two directions (Fig. 4c); AT2 cell changes aligned with the LUAD description, echoing the AT2-related transcriptional lineage in human LUAD and the attenuation of alveolar identity during tumor progression.^26^

Interchangeable disease directions also screen for disease-associated candidate genes in healthy cells. In healthy alveolar macrophages primarily from the Human Lung Cell Atlas (HLCA),^27^ we upregulated single genes *in silico* and read responses against LUAD, pulmonary sarcoidosis, and COVID-19 descriptions (Fig. 4d): CD163 and LIPA were positive in all three directions; FABP4, CXCL16, and C1QB were positive in LUAD and negative in COVID-19, whereas PKM and CD68 showed the opposite pattern; and SLC11A1 and VSIG4 favored pulmonary sarcoidosis, showing that candidate gene priority also depends on the disease of interest. These results corroborate known disease biology: FABP4⁺ alveolar macrophages are depleted in bronchoalveolar lavage fluid from patients with severe COVID-19,^28^ whereas glycolytic genes including PKM are upregulated in alveolar macrophages from COVID-19 decedents,^29^ matching their opposite COVID-19 responses; and SLC11A1 polymorphisms are associated with sarcoidosis susceptibility across multiple populations.^30,31^

To further resolve bidirectional dose responses of multigene programs within diseased cells, we applied 11 bidirectional expression adjustment levels to 85 genes in LUAD epithelial cells from the Lung Cancer Atlas (LuCA)^32^ and read responses along the LUAD direction. Response profiles clustered by direction and amplitude into five groups corresponding to surfactant/MHC-II, secretory/immune, near-inert, keratin/metabolic, and invasion/EMT programs (represented by FOLR1, IFITM2, CALM2, NDUFS5, and ANXA2, respectively; Fig. 4e): upregulating malignancy and invasion genes (SOX4, SEC61G, ANXA2, S100A10) shifted representations toward the LUAD description and downregulating them shifted representations away, whereas alveolar differentiation genes (NAPSA, SFTPB, FOLR1) and antigen-presentation genes (CD74, HLA-DRA) acted in the opposite direction. Consistent with tumor biology, SEC61G promotes LUAD proliferation, invasion, and metastasis,^33^ NAPSA and SFTPB belong to the AT2 surfactant program,^34^ loss of the alveolar differentiation master regulator NKX2-1 characterizes high-grade poorly differentiated LUAD in mouse models,^35^ and tumor-intrinsic MHC-II expression shapes the LUAD immune microenvironment,^36^ decomposing the LUAD state into interpretable components of reduced differentiation, weakened antigen presentation, and enhanced invasion.

Two-gene perturbations further test interactions within and across programs. Co-downregulating gene pairs in the alveolar differentiation program and co-upregulating pairs in malignancy/invasion programs yielded positive non-additive effects in all combinations of the former and most of the latter (Fig. 4f and Supplementary Fig. S5e). Joint weakening of alveolar genes such as NAPSA–SFTPB and joint strengthening of malignancy genes such as SEC61G–S100A10 converged on the same disease state from opposite directions (Supplementary Fig. S5a,b), consistent with loss of lineage identity during LUAD evolution,^37^ while Nkx2-1 haploinsufficiency likewise synergizes with oncogenic Kras G12D to promote tumor progression in mice.^38^ Combinatorial genetic perturbations can depart from additive expectations,^39^ and their direction also depends on functional context: co-downregulation within the antigen-presentation program (such as CD74–HLA-DRA) consistently produced positive non-additive effects, whereas co-upregulation between malignancy/invasion and immune programs almost uniformly showed buffering (such as SOX4–HLA-DRA; Supplementary Fig. S5c–e), consistent with reports that reducing tumor MHC-II expression decreases T-cell infiltration^36^ and that SOX4 acts as a tumor-intrinsic factor conferring resistance to T-cell killing.^40^ Fixing the NAPSA–SFTPB co-downregulated state while switching disease descriptions shifted this state toward LUAD and lung squamous cell carcinoma (LUSC) and away from COVID-19 and idiopathic pulmonary fibrosis (IPF; Fig. 4g and Supplementary Note 3), showing that the same combinatorial change carries different meanings under different disease questions. Natural language thus extends from describing cells to serving as a reference for exploring gene–disease relationships.

### LingoCell resolves cell identities and identity-conditioned niches at single-cell resolution in spatial transcriptomics

Through measurement adaptation and spatial self-supervised pretraining, LingoCell outputs both an identity embedding and a niche embedding for each segmented spatial cell (Figs. 1e,f and 5). Because measurement adaptation preserves the prototype geometry learned during single-cell pretraining, LingoCell directly matches the identity embeddings of Visium HD segmented cells to cell-type prototypes, resolving 20, 27, and 27 cell types in kidney, normal colon, and lung cancer sections, respectively, without a matched single-cell reference (Fig. 5a and Supplementary Figs. S6a and S7a); mapped back onto H&E, all populations followed tissue morphology and were supported by marker genes (Supplementary Figs. S6b, S7b, and S8). In kidney, proximal tubules (CUBN, LRP2), loops of Henle (SLC12A1, UMOD), distal and connecting tubules (SLC12A3, TRPM6), collecting ducts (AQP2, SLC4A1), and podocytes (NPHS2, PODXL) aligned along the nephron and separated in the identity embedding, with in situ expression of SLC12A1, SLC12A3, AQP2, and NPHS2 localizing to the corresponding annotated structures in the zoomed-in field (Fig. 5a).

**Fig. 5.**
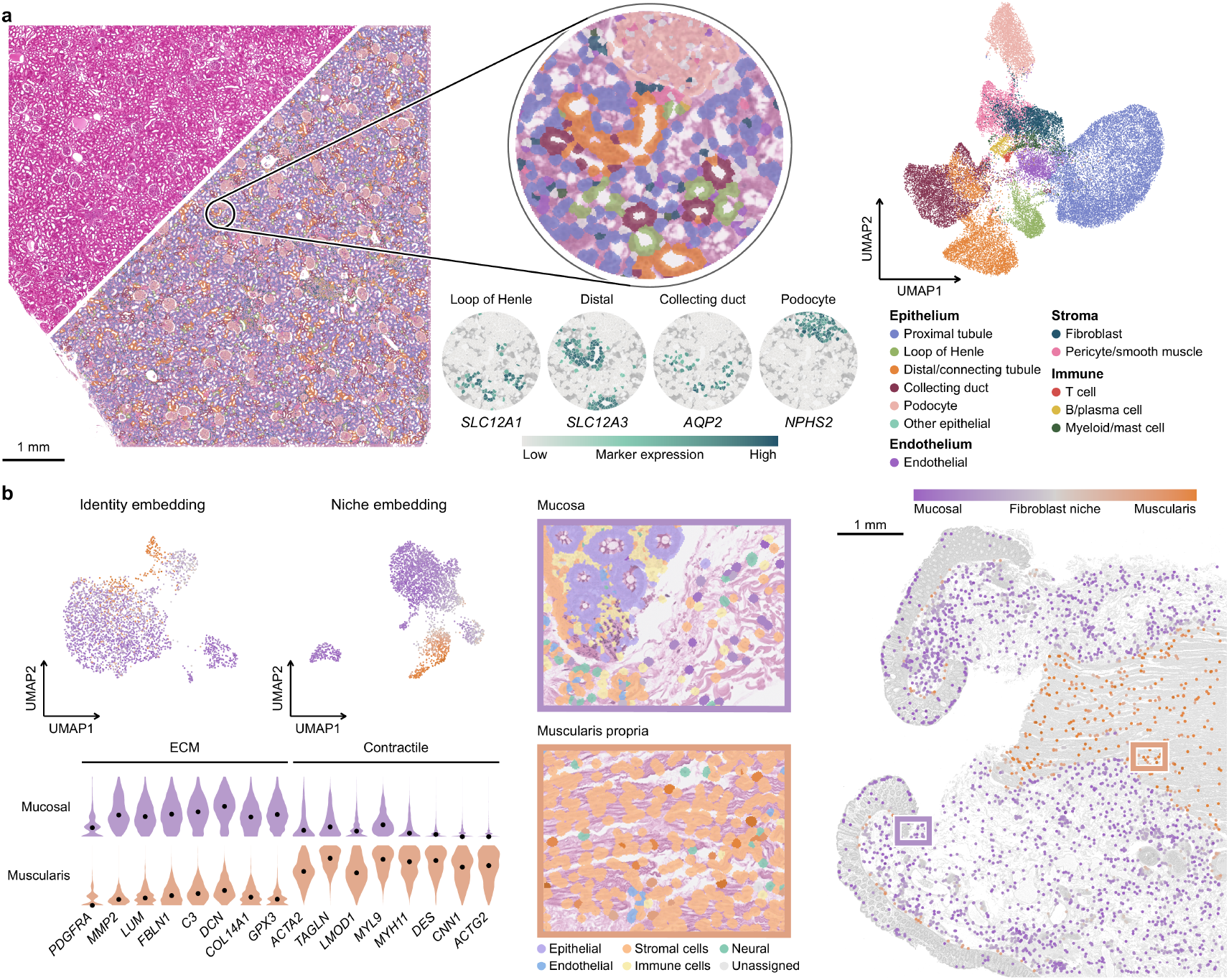
**LingoCell resolves cell identities and identity-related local niches at single-cell resolution**. **a,** Cell identity annotation in a kidney section: the whole section with H&E and annotations (left), segmented cell outlines overlaid on H&E in the circled field of view (upper middle), in situ expression of SLC12A1 (loop of Henle), SLC12A3 (distal tubule), AQP2 (collecting duct), and NPHS2 (podocytes) in the same field (lower middle), and a UMAP of identity embeddings (right); colored by cell class. Scale bar, 1 mm. **b,** Identities and niches of normal colon fibroblasts: UMAPs of the same fibroblasts in identity and niche embeddings (upper left), expression of extracellular matrix and contraction genes in neighbors of mucosa-associated and muscularis-associated fibroblasts (lower left), enlarged views of the mucosa and muscularis propria with fibroblasts colored by niche coordinate and other cells by lineage (middle), and the distribution of fibroblasts across the whole section colored by niche coordinate (right; rectangles mark the enlarged views). Scale bar, 1 mm.

Identity annotation answers what a cell is, but not what environment it occupies. Unlike principal component analysis (PCA), which summarizes variation in a cell’s own expression, the niche embedding characterizes the surrounding environment referenced to the cell’s own identity, subdividing populations by microenvironment within the same cell type. In normal colon (Fig. 5b), the same fibroblasts intermingled in the identity embedding but separated in the niche embedding into mucosal and muscularis groups: mucosal fibroblasts lay among crypt epithelium, immune cells, and loose stroma with neighbors enriched for extracellular matrix genes (DCN, LUM, C3, COL14A1), whereas muscularis fibroblasts lay between smooth muscle bundles with neighbors enriched for contractile genes (DES, MYH11, TAGLN, ACTA2), their difference defining a contractile–extracellular matrix axis (Supplementary Note 4). This distinction persisted among cells with nearly identical neighbor cell-type compositions (Supplementary Fig. S6c,d), showing that the niche embedding captures continuous neighbor molecular states beyond discrete neighbor categories and provides a representation basis for analyzing neighborhood gene programs.

### LingoCell resolves observer-dependent stromal programs in colorectal cancer through spatial *in silico* perturbation

To test whether the model can resolve cell identities and microenvironmental interactions in highly heterogeneous tumor tissue, we applied LingoCell to the colorectal cancer (CRC) Visium HD data of de Oliveira et al.^7^ Whereas the original study annotated 8 μm bins with RCTD,^41^ LingoCell directly annotated segmented cells in section P5, yielding consistent tissue partitions supported by the identity UMAP and canonical marker expression (Fig. 6a,b and Supplementary Fig. S9a). Agreement on major cell types was 88.0% and reached 97.6% in homogeneous single-type RCTD regions, with differences concentrated at tissue boundaries containing mixed cell types (Supplementary Fig. S9b); in these mixed units, LingoCell recovered hundreds of T cells within RCTD-labeled tumor regions that expressed T-cell markers while surrounding tumor cells barely did (Fig. 6c and Supplementary Fig. S9c).

**Fig. 6.**
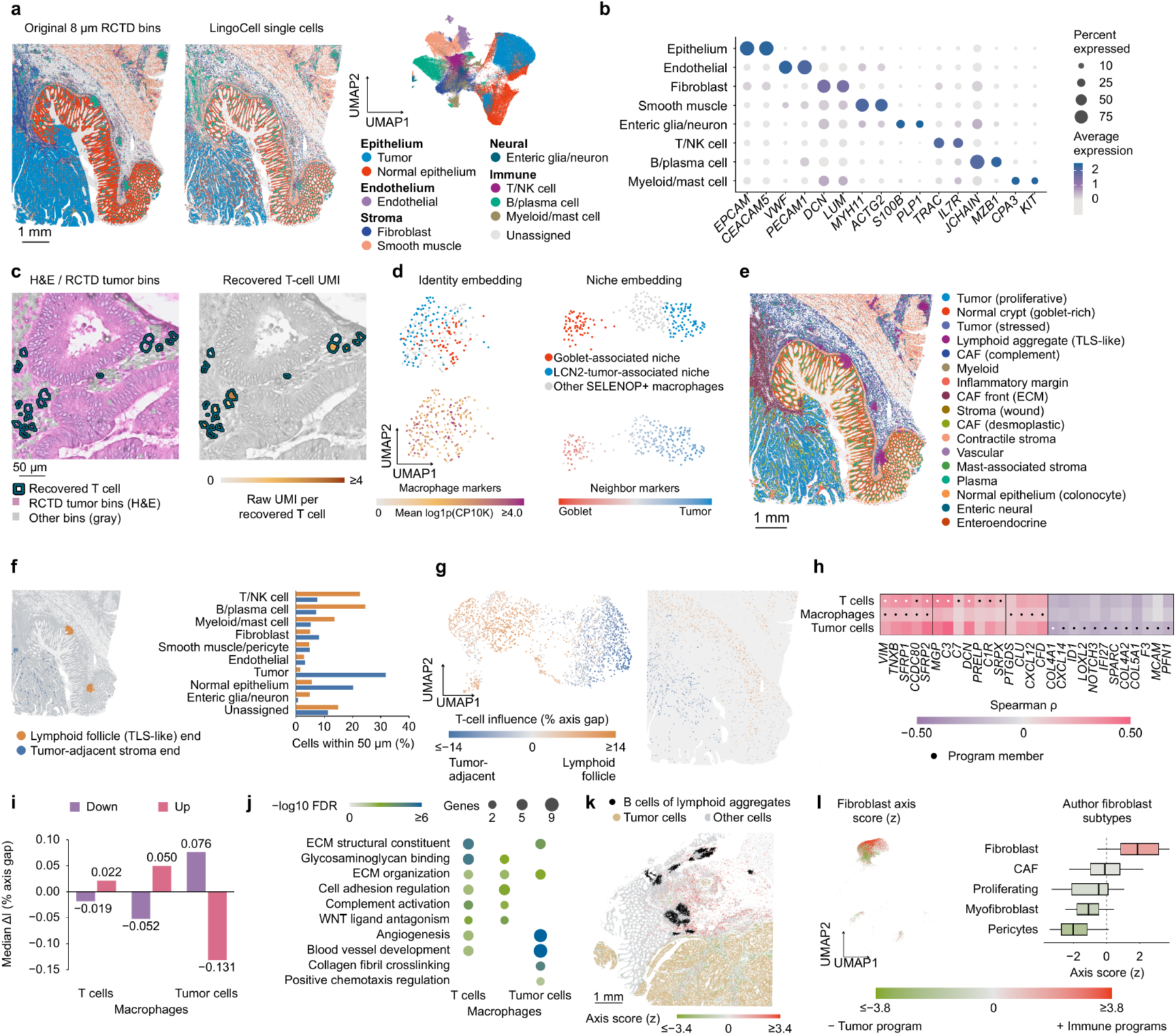
Spatial *in silico* perturbation resolves observer-dependent stromal programs in CRC. a, RCTD annotations of 8 μm bins from the original study (left), LingoCell annotations of segmented cells (middle), and an identity embedding UMAP (right) in section P5, colored by cell class. Scale bar, 1 mm. **b,** Dot plot of canonical marker gene expression across cell classes (dot size, detection rate; color, standardized mean expression). **c,** Example field of T cells recovered by LingoCell in a region annotated as tumor by RCTD: H&E with outlines of recovered cells (left) and T-cell marker gene expression (right). Scale bar, 50 μm. **d,** UMAPs of the same SELENOP⁺ macrophages in identity and niche embeddings, colored by goblet-associated and LCN2 tumor-associated neighborhoods and by marker expression in the macrophages or their neighbors. Three outlying cells are omitted from the identity-embedding displays. **e,** Functional niches grouped from niche embedding clusters. Scale bar, 1 mm. **f,** Two reference environments for spatial perturbation: anchor T cells at the two ends of the reference axis (orange, lymphoid follicle (TLS-like) end; blue, tumor-adjacent stroma end) and cell composition within 50 μm. **g,** Influence of fibroblast donors from the T-cell perspective in identity embedding UMAP (left) and tissue coordinates (right). **h,** Correlations between program gene expression in fibroblast donors and influence from T-cell, macrophage, and tumor-cell perspectives (dots mark members of each observer program). **i,** Median ΔI of fibroblast donors after bidirectional program editing (purple, downregulation; pink, upregulation). **j,** Functional enrichment of the three programs in external CRC fibroblasts (dot size, overlapping gene count; color, −log₁₀FDR). **k,** Axis scores of fibroblasts in section P1 (black, B cells in the lymphoid aggregate; sand, tumor cells; scale bar, 1 mm). **l,** Axis scores of fibroblasts in the P1–P5 single-cell atlas from the same study: original UMAP (left) and box plots grouped by fibroblast subpopulations (right). CRC, colorectal cancer; TLS, tertiary lymphoid structure; FDR, false discovery rate.

Niche embeddings further distinguished tissue contexts sharing the same cell identity. Whereas the original study combined Visium HD and Xenium to identify SELENOP/STAB1 macrophages near REG1A⁺LCN2⁺ tumor cells and goblet cells,^7^ LingoCell reproduced both environments from Visium HD alone: both macrophage groups retained SELENOP–STAB1–C1Q features and clustered together in the identity embedding but separated clearly in the niche embedding, with neighbors enriched for goblet-cell markers (MUC2, ITLN1, TFF3) and tumor markers (LCN2, CEACAM5, CEACAM6), respectively (Fig. 6d).

Across the whole section, niche embedding clusters annotated and merged by cell composition, marker genes, and pathway activity yielded 17 functional niches spanning tumor states, lymphoid aggregates (TLS-like), multiple cancer-associated fibroblast (CAF) zones, and vascular and contractile stroma (Fig. 6e and Supplementary Fig. S10), with lymphoid aggregates forming sharply bounded niches whereas corresponding scGPT-spatial and GraphST clusters intermingled within the aggregate or spread outward (Supplementary Fig. S11). Tertiary lymphoid structures (TLS) recruit and organize T cells in tumors, correlate in density with CRC T-cell infiltration and prognosis,^42^ and are supported by stromal cells including CCL19⁺ fibroblasts and fibroblastic reticular cells (FRC).^43,44^ Because the P5 lymphoid follicle comprised a B-cell follicle, T cells, and CCL19⁺ fibroblasts, we used its T cells as observers to interrogate the environmental information carried by neighboring fibroblasts.

We selected T/B-cell-rich lymphoid follicle (TLS-like) regions and lymphocyte-poor, malignant-epithelium-rich tumor-adjacent stroma in P5 as reference environments: both contained fibroblasts (Fig. 6f), the neighbor type eliciting the largest variance in model response and lying immediately adjacent to follicular T cells (Supplementary Fig. S12). However, because the two regions differed simultaneously in cell composition, spatial structure, and molecular state, direct comparison could not determine which stromal features shaped the T-cell environment. Spatial *in silico* perturbation converts this question into a direct model query: fixing T cells and their surrounding context while replacing neighborhood fibroblasts in turn with each real donor across the section, we measured donor influence by the median shift of the observer’s niche embedding along the tumor-adjacent stroma–lymphoid follicle reference axis (positive values indicating a more lymphoid-follicle-like environment). Because all donors are compared within the same local context, influence directly reflects state differences within fibroblasts: it formed a continuous gradient in the donor identity embedding, and positive and negative donors occupied distinct tissue regions when mapped back onto the section (Fig. 6g).

We identified fibroblast genes consistently correlated with influence across two non-overlapping spatial groups and combined co-expression with an external CRC single-cell fibroblast reference to construct gene programs (Methods).^45^ The positive T-cell program comprised MGP, C3, VIM, TNXB, SFRP1, C7, DCN, CCDC80, SFRP2, PRELP, C1R, and SRPX, enriched for reticular extracellular matrix organization and complement activation (Fig. 6h,j). Downregulating this program in real donors shifted T-cell environmental readouts away from the lymphoid follicle end (q = 3.16 × 10⁻⁴), whereas upregulation shifted them toward it (q = 8.60 × 10⁻⁶; Fig. 6i and Supplementary Table S6a). The program overlapped with a CCL19-marked stromal state in the external atlas (Supplementary Fig. S13g and Supplementary Table S6f);^45^ CCL19 was detected more frequently near the P5 follicle end but was too sparse to enter candidates (Supplementary Fig. S13a,b), indicating that the model reads a more stably expressed composite signal reflecting a CCL19-associated lymphoid-type stromal state. We therefore define it as a CCL19-associated reticular extracellular matrix–complement fibroblast program, consistent with reports that CCL19⁺ fibroblasts and FRC support T-cell organization.^43,44^

Changing the observer altered the genes prioritized from the same fibroblast state. With macrophages as observers, the positive program (SFRP2, PTGDS, SFRP1, CCDC80, TNXB, VIM, CLU, CXCL12, CFD) retained the shared SFRP1–SFRP2–CCDC80–TNXB–VIM backbone while adding CXCL12, CFD, and CLU to highlight chemokine sensing and complement processing (Fig. 6h,j and Supplementary Table S6f), where CXCL12 can form a stromal–myeloid signaling interface via CXCR4.^46^ With tumor cells at the opposite end of the reference axis as observers, the model read a distinct program (COL4A1, CXCL14, ID1, LOXL2, NOTCH3, IFI27, SPARC, COL4A2, COL5A1, F3, MCAM, PFN1) dominated by blood vessel development, angiogenesis, and collagen fibril crosslinking, matching perivascular stromal states in the external atlas (Fig. 6h,j and Supplementary Table S6f,g)^45^ and consistent with reports that CXCL14⁺ fibroblasts line tumor epithelium in CRC and that high LOXL2 expression in colon CAFs correlates with poor prognosis.^47,48^ Bidirectional editing shifted both observers in the expected directions (macrophage downregulation and upregulation adjusted P = 3.06 × 10⁻⁵⁸ and 2.80 × 10⁻³⁸; tumor cells < 10⁻⁸⁴ in both directions; Fig. 6i and Supplementary Table S6a).

Across all three observers, positive T-cell and macrophage program genes correlated positively with influence, whereas negative tumor-cell program genes correlated negatively (Fig. 6h): rather than identifying three unrelated fibroblast subtypes, the model recovered a shared stromal axis extending from a lymphoid-follicle-associated reticular and complement stromal state to a vascular, basement-membrane, and matrix-remodeling state, assigning different priorities to genes along this axis according to observer identity. Defining an axis score by the standardized score difference between immune-observer and tumor-observer programs, fibroblasts around lymphoid aggregates in section P1 of the same study favored the immune end whereas inter-tumoral stroma favored the tumor end (Fig. 6k, Supplementary Figs. S13c and S14a, and Supplementary Table S6b,c); in the matched single-cell atlas, the two ends corresponded to the original study’s Fibroblast subpopulation and to pericyte and myofibroblast subpopulations (Fig. 6l and Supplementary Fig. S14b); and in the independent Valdeolivas standard Visium cohort, both immune-observer programs scored higher in lymphoid-aggregate-adjacent stroma across most subjects (Supplementary Fig. S15, Supplementary Table S6d,e, and Supplementary Note 5).^49^ Whereas analyses based on neighborhood composition, colocalization, or ligand–receptor pairing mainly describe what is present in an environment, observer-conditioned spatial *in silico* perturbation further addresses which environmental components convey information to a specified immune or tumor cell.

## Discussion

LingoCell unifies editable molecular inputs with interpretable biological references within a single foundation model: the single-cell stage supports precise editing through joint encoding of gene identity and expression magnitude while providing disease-concept readouts through natural-language alignment, and the spatial stage uses a cell’s own identity as the query reference for reading neighborhood states. Single-cell and in situ tissue analyses thus follow the same paradigm, modifying molecular inputs within a defined cellular context and interpreting responses along biologically intelligible directions.

Semantic perturbation requires that expression changes both enter the model and acquire biological meaning: expression models such as scGPT are magnitude-sensitive but lack language references,^2^ whereas alignment models such as LangCell have language references but are restricted by rank-based encoding to deleting or fronting genes.^1,4^ LingoCell unifies both in a single encoder so that genes can be adjusted stepwise across ordered levels and read directly along disease-language directions. Expression changes and language references can thus be controlled independently: fixing the disease reference compares direction, amplitude, and non-additivity across single-gene and combinatorial edits, whereas fixing the edited state while switching language references re-examines the same molecular change across diseases without retraining classifiers, making language an analytical coordinate adaptable to the research question.

Transferring to spatial data is challenged by sparse capture altering how the same cell appears, such that direct continual training can misread measurement sparsity as identity differences. LingoCell’s measurement adaptation training teaches the model to distinguish lower capture depth from a different cell identity before learning spatial context; the preserved cell-type knowledge supports reference-free spatial annotation and provides a stable reference for niche comparisons within the same cell type.

Previous methods mostly define niches from the environment itself, aggregating neighborhood information without conditioning on the central cell’s identity.^11,12^ LingoCell instead starts from who is observing, using the central cell’s own identity as a query to generate a niche embedding complementary to its identity embedding. The environment thus comprises continuous neighbor molecular states rather than only discrete neighbor categories, distinguishing microenvironments with similar neighbor compositions; the same neighbor state is read as different environments under different observer identities, matching the biological intuition that the same stroma carries different meanings for T cells, macrophages, and tumor cells.

This structure allows neighbor states to be individually replaced or edited and re-evaluated through the environmental computation, localizing environmental readouts to candidate gene programs and testing their effect directions. Because conclusions derive from changes in full model outputs upon input modification rather than single-layer attention weights, the resulting programs have explicit observers, directions, and comparison contexts, providing prioritized leads for validating immune and peritumoral stromal interactions.

In spatial cells with limited molecular coverage, fine-grained state resolution remains bounded by observable transcripts, and expanding pretraining across tissues, platforms, and language descriptions should broaden applicability; combining expression–semantic and expression–environment relationships from *in silico* perturbation with Perturb-seq and spatial functional experiments will help connect predictions to biological responses, while richer text and morphological inputs can be naturally integrated. Overall, LingoCell enables the meaning of genes and their combinations to be interpreted across both disease semantics and tissue environments, offering a unified approach to prioritizing targets and generating testable hypotheses from cell atlases and tissue sections.

## Methods

### Data sources and preprocessing

#### Data sources

Expression matrices and cell annotations for single-cell pretraining were obtained from CZ CELLxGENE Discover and combined with paired cell descriptions to construct the expression–language corpus (Supplementary Table S1a).^13^ Spatial pretraining used publicly released Visium HD data from 10x Genomics (Supplementary Table S1b), with inputs including cell-associated expression, spatial coordinates, and registered tissue image features. Single-cell downstream analyses used PBMC10K, Pancreas, Human Immune, and Tabula Sapiens to evaluate cell representations and annotation, and the MacParland human liver atlas and lung atlases for description generation, semantic queries, and *in silico* perturbation (Supplementary Table S1c). Spatial applications used kidney, tumor-adjacent normal colon, and lung cancer data to resolve cell identity and local environments, together with the CRC Visium HD data of de Oliveira et al.,^7^ with the P5 section used for identity, niche, and *in silico* perturbation analyses and the P1 section, the matched single-cell atlas, and data from independent studies used for program expression and spatial localization analyses (Supplementary Table S1d).

#### Single-cell data preprocessing

The raw expression matrix is denoted *X* ∈ ℝ*^N^*^×*G*^ (*N* cells, *G* genes, with *x_i_*_g_ the observed expression of gene *g* in cell *i*). After gene identifiers are mapped to a unified vocabulary, each cell undergoes total-count normalization, log1p transformation, and within-cell ordered binning to obtain discrete expression levels *v_i_*_g_:

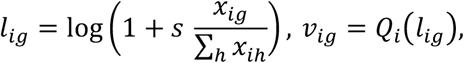

where *s* is the total-count normalization scale, *l_i_*_g_ is the transformed expression, and *Q_i_* is the binning map for cell *i* (51 expression levels), retaining up to 1,200 nonzero-expressed genes per cell.

#### Spatial transcriptomic data preprocessing

Visium HD data were processed using 10x Genomics Space Ranger (v4.1). Nuclei and cells were segmented using registered H&E images, and spatial expression signals were associated with segmented cells to obtain cell-associated expression matrices and spatial coordinates.

### Single-cell expression encoding and input representation

#### Expression input and gene representations

The expression encoder receives two components: gene tokens representing gene identity and discrete expression levels representing the expression of the corresponding genes. Let *G_i_* denote the set of genes actually entering the model for cell *i* and *v_i_*_g_ the discrete expression level of gene *g*. The cell input is represented as:

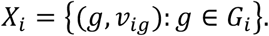

Gene identities are converted to vectors 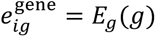 through a unified vocabulary and a learnable gene embedding mapping *E*_g_, and discrete expression levels *v_i_*_g_ are mapped to expression value vectors 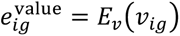) through a multilayer perceptron *E_v_*; the two are summed to obtain the gene representation supplied to the Transformer:

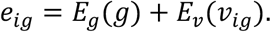

#### Expression value encoding

The discrete expression level *v_i_*_g_ is mapped to an expression value vector through a multilayer perceptron *E_v_*:

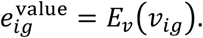

A CLS token is added at the beginning of the input sequence, and its corresponding Transformer output is used to obtain the identity embedding of the cell. Padding positions do not participate in valid attention calculations or the expression recovery objective.

#### Gene expression encoder

The expression encoder uses a Transformer architecture that jointly encodes gene identity and expression value (12 layers, 8 attention heads per layer, hidden dimension 512), similar to scGPT, and is trained from scratch.^2^ For cell *i*, the CLS token input embedding *e*_CLS_ and the joint representations *e_i_*_g_ of the input genes are passed to the expression Transformer *F_θ_* to obtain the output sequence:

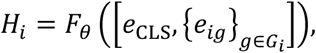

where representations at gene positions are used for expression recovery and conditional language generation.

#### Identity embedding

The output at the CLS position, *H_i_*_,CLS_, is sequentially subjected to layer normalization (LN) and L2 normalization (norm) to obtain the identity embedding used for cell annotation, clustering, cross-batch integration, and semantic perturbation:

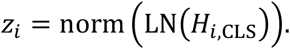

### Text encoding and language generation

#### Text encoder

The paired cell description *T_i_* is encoded by PubMedBERT (*B_φ_*, 12 Transformer layers with a hidden dimension of 768) initialized with pretrained weights.^50^ Its CLS output is mapped by a projection network *A_t_* trained from scratch to the same 512 dimensions as the identity embedding and normalized to obtain the text representation:

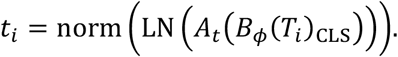

#### Expression–text alignment

For each cell, its paired description and descriptions with the same cell identity form the positive set *P*(*i*), and *C*(*i*) denotes the set of valid candidate texts. The expression-to-text contrastive loss is:

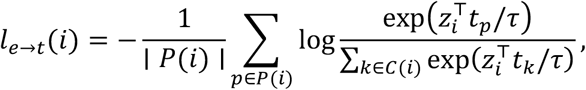

where *t_p_* and *t_k_* are the text representations of positive text *p* and candidate text *k*, respectively, and *τ* is the similarity scaling parameter. Swapping expression and text yields the reverse loss, and the two are averaged to obtain the alignment loss *L*_align_.

#### Language decoder

The language decoder *p*_A_ is based on TinyLlama (22 Transformer layers with a hidden dimension of 2,048) initialized with pretrained weights,^51^ with cross-attention added to its last 12 layers to read cell expression information. The expression encoder output sequence *H_i_* is converted by an adapter *A*_g_ into the conditional input for the decoder, which generates the reference description *w_i_*_,1:*K*_*_i_* (valid length *K_i_*) token by token under the description instruction *q*. The language generation loss is:

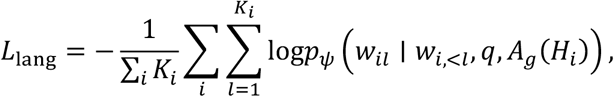

where *w_i_*_,<*l*_ denotes the reference text preceding the *l*th token.

### Single-cell expression–language pretraining

Single-cell pretraining proceeds in two stages: the first stage takes cell expression, paired text descriptions, and cell-type labels as inputs to jointly learn the expression encoder, text encoder, and cell-type prototypes; the second stage freezes the expression encoder and trains the adapter and language decoder for expression-conditioned language generation.

#### Expression recovery

Expression levels are masked at 40% of positions among genes with nonzero expression and predicted using contextual representations at gene positions, text conditions, and identity embeddings combined with target gene identities, respectively. The expression recovery loss *L*_expr_ measures the difference between predicted and true expression levels, allowing the encoder to retain gene expression information.

#### Identity prototype constraints

Cell-type prototypes covering 785 cell types are constructed on the expression and text sides separately, and known type labels constrain the similarity between identity embeddings and their corresponding prototypes. The identity prototype loss *L*_proto_ provides a shared identity reference for cells of the same type.

#### Two-stage training objectives

In the first stage, the joint representation learning loss is a weighted sum of three terms (*λ* denotes the weights of the loss terms):

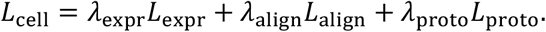

In the second stage, with the expression encoder frozen, the adapter *A*_g_ and language decoder *p_ψ_* are trained by minimizing the language generation loss *L*_lang_ conditioned on the cell expression sequence and description instruction *w_i_*_,1:*Ki*_.

### Measurement adaptation training for spatial transcriptomics

To bridge the differences in capture depth and sparsity between single-cell and spatial transcriptomics, LingoCell introduces measurement adaptation training between single-cell pretraining and spatial pretraining, adapting the expression encoder to spatial measurement conditions while retaining existing cell identity and semantic references.

#### Observation transformation

An observation transformation *T* targeting the spatial measurement distribution is applied to the reference expression input *X_i_* of each cell to obtain expression input under spatial measurement conditions:

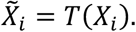

#### Adaptation training objectives

Training jointly optimizes three objectives: an expression recovery loss *L*_obs_ that predicts expression in the transformed observation, an identity consistency term that constrains the identity embeddings of the same cell under *X*^c^*_i_* and *X_i_* to remain consistent, and a semantic preservation term *L*_sem_ that retains existing expression–text relationships and cell-type prototype references.

### Identity-conditioned spatial encoding

#### Neighborhood input

The spatial model combines central-cell expression, neighbor expression, spatial distances, and tissue image information to learn identity and niche embeddings. Expression of the central cell and its 20 spatially nearest neighbors is processed by the adapted expression encoder, and neighbor identity embeddings are combined with encodings of the distances from the central cell to its neighbors.

#### Tissue image feature extraction

Registered H&E image patches centered on each cell are divided into four subregions, and the frozen pretrained pathology vision model MUSK extracts 1,024-dimensional image embeddings from each subregion,^52^ which are mapped through an adapter and supplied as image inputs to the spatial model.

#### Spatial information aggregation

The central cell’s own gene expression is encoded by the adapted expression encoder to obtain the identity embedding ℎ*_i_* used to identify cell type (denoted *u_i_* = ℎ*_i_*/∥ ℎ*_i_* ∥ _2_ after L2 normalization). Using ℎ*_i_* as a query, the spatial attention module integrates the identity embeddings of neighboring cells, spatial distances, and tissue image features through 8-head cross-attention to obtain environmental information *s_i_*. After mapping, the component along the central cell’s identity direction is removed to obtain the niche embedding describing the local environment:

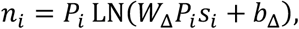

where 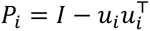 is the orthogonal projection operator for the identity direction, *W*_Δ_ and *b*_Δ_ are learnable parameters, and LN denotes layer normalization.

### Spatial pretraining

#### Spatially conditioned expression recovery

During training, expression levels of some genes in the central cell are masked, and the model combines the resulting gene representations with aggregated spatial context to predict expression levels at the masked positions. The mean squared error between predicted and measured expression levels gives the spatial expression recovery loss *L*_spatial expr_, incorporating neighbor and tissue image information into the prediction of central-cell expression.

#### Neighborhood summary prediction

Cell-type prototypes from single-cell pretraining are used to compute soft identity assignments for neighbors, and AUCell activities of pathway gene sets represent their molecular activities.^53^ Weighted aggregation yields summaries of neighborhood cell composition and molecular activity, which the model predicts from the central cell’s niche embedding, with the niche loss *L*_niche_ measuring the differences between predictions and summary targets.

#### Training implementation

LingoCell was implemented in PyTorch and trained sequentially through single-cell expression–language pretraining, expression-conditioned language decoder training, spatial measurement adaptation, and spatial self-supervised pretraining on 16 NVIDIA H100 GPUs; all downstream analyses were performed by direct inference with the frozen pretrained models.

### Single-cell representation evaluation and description generation

#### Cell clustering

A nearest-neighbor graph is constructed from identity embeddings extracted by the frozen expression encoder for Leiden clustering.^54^ For comparison with reference cell types, the partition with the highest NMI is selected from a common set of candidate resolutions, and ARI is calculated for that partition:

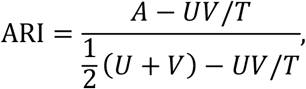

where *A* is the number of cell pairs belonging to both the same reference type and the same predicted cluster, *U* and *V* are the numbers of cell pairs belonging to the same reference type and the same predicted cluster, respectively, and *T* is the total number of cell pairs.

#### Cross-batch integration

Cells from different batches are passed through the same frozen expression encoder, a nearest-neighbor graph is constructed from the combined identity embeddings, and biological conservation and batch mixing are evaluated using scIB metrics:^15^

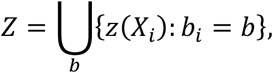

where *z*(*X_i_*) is the identity embedding of cell *i* and *b_i_* is its batch label. LingoCell does not use batch labels of the target data to generate representations. Harmony, scVI, and Scanorama are compared as integration methods that use batch information.^17–19^

#### Zero-shot cell annotation using text references

Without training or fine-tuning on the target data, cosine similarities are calculated directly between the identity embedding *z_i_* of each cell to be annotated and the text representations *t_c_* of candidate cell-type descriptions, and the candidate with the highest similarity is assigned as the predicted type *ŷ_i_*:

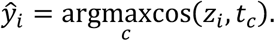

#### Cell description generation and evaluation

Gene representations of cells are passed through the adapter to the language decoder to generate natural-language descriptions. Generated descriptions are compared with reference descriptions using BERTScore based on RoBERTa and BioBERT for semantic similarity,^55–57^ BLEU and ROUGE for lexical matching,^58,59^ and weighted F1 scores for cell-type, tissue, and disease information extracted from the generated text:

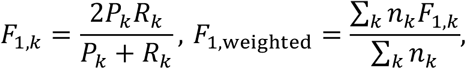

where *P_k_* and *R_k_* are the precision and recall of information extraction for class *k*, respectively, and *n_k_* is the number of cells in that class in the reference descriptions. In the human liver single-cell data, zonation descriptions generated for hepatocytes are further compared with the expression distributions of zonation-related genes.

#### Gene association analysis of generated descriptions

Cross-attention to the set of analyzed input genes *G*_i_* is extracted when the language decoder generates the hepatocyte zonal identity phrase *w_i_*. With *R_i_*_g_ denoting the relative attention rank of gene *g* within cell *i* (higher attention corresponds to a larger value), the difference in mean relative rank between the target phrase group *w* and other phrase groups is calculated:

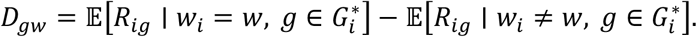

A larger *D*_gw_ indicates a stronger relative association between gene *g* and target phrase *w*.

### Single-cell semantic querying and *in silico* perturbation

#### Natural-language queries and disease-state analysis

With *t*_1_ and *t*_O_ denoting the normalized text representations of the target biological process or disease description and its reference description (homeostatic descriptions for process queries, and normal descriptions matched for cell type and tissue context for disease queries), the semantic query score is the projection of the identity embedding *z_i_* onto their difference:

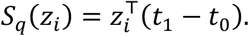

### Single-gene *in silico* perturbation

With the expression encoder frozen, an expression level adjustment *δ* is applied to gene *g* (denoted ℰ_g,ẟ_(*X_i_*)), the cell is re-encoded, and semantic query scores before and after perturbation are compared, giving the semantic shift ΔF in Fig. 1d:

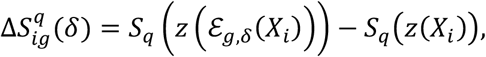

where *z*(⋅) is the identity embedding output by the expression encoder. Single-gene upregulation screens and upregulation response profiles are corrected for the common response within the same cell, whereas downregulation response profiles use raw responses.

#### Two-gene perturbation and combinatorial responses

In cells where both genes *a* and *b* enter the model input, semantic query scores are calculated for the unperturbed state (*z_i_*^0^), the individual perturbations (*z_i_^a^*,*z_i_^b^*), and the joint perturbation (*z_i_^ab^*), and the non-additive effect of the combination is defined as:

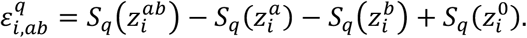

The sign of the non-additive effect indicates enhancement or attenuation along the query direction relative to the sum of the two individual responses.

### Spatial cell annotation and niche analysis

#### Spatial cell identity annotation

According to the tissue of origin of each spatial sample, cell types from the corresponding normal tissue are selected from the single-cell pretraining atlas, retaining immune, stromal, vascular, and other types to form the candidate type set *C_T_*. The identity embedding ℎ*_i_* of each spatial cell is compared with the expression-side prototypes *p_c_* of the candidate types, the type with the highest cosine similarity is assigned as the initial annotation, and the annotations are checked against marker gene expression and spatial location (Fig. 5a and Supplementary Figs. S6–S8):

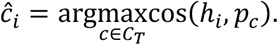

For colorectal cancer sections, malignancy-related prototypes do not participate in initial type assignment. The model first identifies epithelial cells, then calculates their highest similarity to malignancy-related prototypes, using the malignancy similarity distribution of immune and stromal cells in the same section as a reference. When the high-similarity peak of epithelial cells clearly exceeds the upper boundary of the reference distribution, a high quantile of the reference distribution is used as the threshold, and epithelial cells above it are labeled as malignant epithelium (Fig. 6).

#### Neighborhood composition matching and molecular-state analysis

To analyze whether niche embeddings contain information beyond neighboring cell-type composition, we selected 15 nearest neighbors for each central cell among normal colon fibroblasts based on spatial coordinates, calculated the proportions of cell types in each neighborhood, performed nearest-neighbor matching between muscularis-associated and mucosa-associated niche groups on these proportions without reusing matched cells, and measured compositional differences by the L1 distance between the mean neighborhood compositions of the two groups. From the mean expression *x_ig_* of gene *g* in the neighborhood of cell *i*, a continuous molecular axis is defined as the difference in mean expression between the contraction-related gene set *G*_con_ and the extracellular matrix-related gene set *G*_mat_:

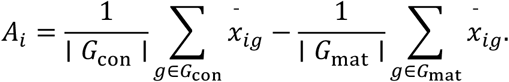

Principal component analysis is performed on the niche embeddings of matched cells, and the first principal component (niche PC1, with its display direction aligned with the molecular axis) serves as a continuous environmental coordinate whose Pearson correlation with the molecular axis is calculated. Linear regression then removes the components of each explained by neighboring cell-type proportions, and the correlation between the residuals evaluates their association after controlling for neighborhood composition.

### Spatial *in silico* perturbation and gene program analysis

#### Tissue reference axes

Tumor-adjacent stroma and lymphoid follicle (TLS-like) regions in colorectal cancer tissue serve as reference environments. The mean niche embeddings *a_c_*^-^ and *a_c_*^+^ of observer type *c* in the two regions are calculated separately, and their difference defines a tissue reference axis on which the coordinate of any niche embedding *n* is:

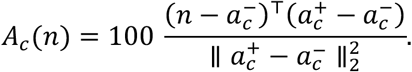

The coordinates of the two reference endpoints are 0 and 100, and a separate reference axis is constructed for each observer type.

#### Neighbor-state replacement and influence calculation

Central cells serve as observers, and the fibroblasts used to replace neighbors serve as donors. Taking T cells as an example, fibroblast donors are selected from the entire section, and each donor’s identity embedding in turn replaces the identity embeddings of four existing fibroblasts around a T cell, with the T cell’s own expression, other neighbors, spatial positions, and tissue images unchanged. After each replacement, the observer’s niche embedding is recalculated, and the influence of donor *d* on observer type *c* is defined as the median of the induced axial change across the observer set *O_c_*:

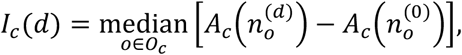

Where *n_o_*^(0)^ and *n_o_*^(*d*)^ are the niche embeddings of observer *o* before replacement and after insertion of donor *d*, respectively. Positive influence indicates that the model’s environmental readout shifts toward the lymphoid follicle endpoint, and negative influence indicates a shift toward the tumor-adjacent stroma endpoint.

#### Influence-associated gene programs

With T cells, macrophages, and tumor cells as observers separately, Spearman correlations are calculated between the expression of each gene in donor fibroblasts and donor influence *I_c_*(*d*). Expression coverage, association directions across spatial groups, gene co-expression, and external fibroblast expression references^45^ are combined to select influence-associated genes into candidate gene programs, and the programs associated with the same fibroblast states are compared across observer contexts.

#### Gene program editing

Genes of a candidate program *G* that actually enter the model are upregulated or downregulated (*σ*) in donor fibroblasts and re-encoded to obtain *d^G^*^,f^. Donors before and after editing (*d*^O^ and *d^G^*^,f^) are placed at the same neighbor positions of the same observer, and influence is compared between the two conditions:

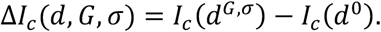

Upregulation and downregulation are summarized separately among donors whose expression was actually modified.

### Statistical analysis

Associations between continuous variables were evaluated using Pearson or Spearman correlation. Nonparametric comparisons across groups were performed using two-sided Mann–Whitney U tests or Kruskal–Wallis tests, and paired comparisons or before-versus-after donor editing comparisons were performed using Wilcoxon signed-rank tests. Overlaps between candidate gene programs and external stromal state marker sets were evaluated using hypergeometric tests, and Gene Ontology (GO) and Reactome pathway enrichment analyses were performed using g:Profiler with false discovery rate (FDR) correction. Multiple hypothesis testing was adjusted using Benjamini–Hochberg (FDR) or Bonferroni correction according to the analysis design. Exact sample sizes, test types, and adjusted statistics for each analysis are reported in the corresponding main text, figure legends, and Supplementary Tables.

## Data availability

All datasets analyzed in this study are publicly available. Single-cell pretraining data were obtained from CZ CELLxGENE Discover (https://cellxgene.cziscience.com/), and spatial pretraining datasets were obtained from the 10x Genomics public Visium HD repository (https://www.10xgenomics.com/datasets). Single-cell benchmark and reference atlas datasets are available through scvi-tools, scIB, Tabula Sapiens, the Human Lung Cell Atlas, and the Lung Cancer Atlas. Additional liver, kidney, brain, and lung single-cell or single-nucleus datasets are available in the Gene Expression Omnibus (GEO) under accession numbers GSE115469, GSE211785, GSE138852, and GSE171524, respectively. Colorectal cancer datasets are available in GEO under accession numbers GSE280318, GSE132465, and GSE144735, and on Zenodo under record 7551712 (https://zenodo.org/records/7551712). Detailed dataset sources and access links are provided in Supplementary Table S1.

## Code availability

The source code and implementation of LingoCell are publicly available on GitHub at https://github.com/y-itao/Lingocell. An interactive web server for online analysis is accessible at https://lingocell.com/.

## Acknowledgements

Computational resources for model inference and device testing were provided by the SHIROKANE supercomputer system at the Human Genome Center, The Institute of Medical Science, The University of Tokyo. Model training and data analyses were performed using the Supermicro ARS-111GL-DNHR-LCC and FUJITSU Server PRIMERGY CX2550 M7 (Miyabi) systems at the Joint Center for Advanced High-Performance Computing (JCAHPC). This work was supported by the Japan Society for the Promotion of Science (JSPS) KAKENHI (grant no. JP26K25585) and the Japan Science and Technology Agency (JST) SPRING program (grant no. JPMJSP2108).

## Author contributions

Y.Y., W.X., and Z.Z. contributed equally to this work. Conceptualization: Y.Y.; methodology: Y.Y. and W.X.; software: Y.Y., W.X., and Z.Z.; formal analysis: Y.Y., W.X., and Z.Z.; investigation: Y.Y., W.X., Z.Z., Y.C., X.Y., and W.G.; data curation: Y.Y., W.X., and Z.Z.; statistical review: Y.Y., Z.Z., Y.C., and J.W.; web server development: W.X.; original manuscript writing: Y.Y. and X.Y.; manuscript review and editing: W.X., Z.Z., X.Y., Y.C., W.G., J.W., and K.N.; supervision: K.N. All authors read and approved the final manuscript.

